# Characterization of a novel peptide mined from the Red Sea Brine Pools and modified to enhance its anticancer activity

**DOI:** 10.1101/2022.01.06.475234

**Authors:** Youssef T. Abdou, Sheri M. Saleeb, Khaled M. A. Abdel-Raouf, Mohamed Allam, Mustafa Adel, Asma Amleh

## Abstract

Peptide-based drugs have emerged as highly selective and potent cancer therapy. Cancer is one of the leading causes of death worldwide. Multiple approaches have been developed towards cancer treatment, including chemotherapy, radiation, and hormonal therapy; however, such procedures’ non-specificity, toxicity, and inefficiency present a hurdle. In this study, we developed a support vector machine (SVM) model to detect the potential anticancer properties of novel peptides through scanning the American University in Cairo Red Sea metagenomics library. Further, we performed in silico studies on a novel 37-mer antimicrobial peptide mined from SVM pipeline analysis. This peptide was further modified to enhance its anticancer activity, analyzed for gene oncology, and subsequently synthesized. The anticancer properties of this 37-mer peptide were evaluated via cellular viability and cell morphology of SNU449, HepG2, SKOV3, and HeLa cells, using MTT assay. Furthermore, we assessed the migration capability of SNU449 and SKOV3 via scratch wound healing assay. Moreover, the targeted selectivity of the peptide for cancerous cells was assessed by testing its hemolytic activity on human erythrocytes. The peptide caused a significant reduction in cellular viability and critically affected the morphology of hepatocellular carcinoma (SNU449 and HepG2), ovarian cancer (SKOV3), and to a limited extent, cervical cancer cell lines (HeLa), in addition to decreasing viability of human fibroblast cell line (1Br-hTERT). Peptide treatment significantly affected the proliferation and migration ability of SNU449 and SKOV3 cells. Annexin V assay was used to evaluate induced cell death upon peptide treatment, attributing programmed cell death (Apoptosis) as the main cause of cell death in SNU449 cells. Finally, we established broad-spectrum antimicrobial properties of the peptide on both gram-positive and gram-negative bacterial strains. Thus, these findings infer the novelty of the peptide as a potential anticancer and antimicrobial agent.

## Introduction

Peptide-based drugs have been gaining more attention as a potential application for anticancer and antimicrobial therapy. They offer greater specificity, fewer off-site side effects, and more potency in comparison to current therapeutic cancer remedies [1]. Biologics (protein-based therapeutics) are large peptide molecules with a typical molecular weight >5 KDa, that are expensive and difficult to produce [2]. Traditional small-molecule drugs with molecular weight <0.5 KDa, manifest non-specific targets, and off-target side effects. Peptide drugs fall within this size gap, ranging between 0.5 and 5 KDa, combining the specificity, potency, and lower toxicity of biologics with the ease of production and metabolic stability of small molecule drugs [2]. Among the classes of peptide drugs are the anticancer peptides (ACPs), which are known for their relatively small sequence (5 to 50 amino acids), and their cationic and amphiphilic properties [2]. Owing to these properties, ACPs specifically target relatively anionic cancer cells while sparing relatively neutral normal mammalian cells [3, 4]. This drug targeting strategy is rarely used in the current cancer therapeutics. Chemotherapeutic agents do not differentiate between normal proliferating cells from cancer cells, thus showing the inability to target indolent or dormant cancers [5]. Cancer cells’ attainment of a chemoresistant phenotype further reduces the therapeutic approach of chemotherapy [6]. Furthermore, current cancer therapy regimens lack in combating multidrug resistance that slowly accumulates in a tumor mass [5].

The anticancer properties of ACPs are established through membranolytic (Direct-acting ACPs) and/or non-membranolytic modes of action (Indirect-acting, programmed cell death) [7]. Anticancer peptides act through the membranolytic mechanisms, disrupt the cellular membrane, mitochondrial membrane, or lysosomal membrane by either the carpet or barrel stave models. Small peptides can aggregate, by hydrophobic interactions, to form a structure through the plasma membrane resembling a traditional ion channel. They can also pass through the plasma membrane and permeate the mitochondrial membrane where they will induce swelling of the mitochondria and release of cytochrome c, subsequently activating caspase 9 and 3. Furthermore, modification of the lysosomal membrane by anticancer peptides can result in acidification of the cytosol. However, the non-membranolytic mechanism of action includes activation of calcium channels resulting in calcium ion influx, augmentation of proteasome activity, inhibition of pro-survival genes, or cell cycle arrest [4, 8]. Thus, ACPs exert a non-systemic-targeted effect when compared to chemotherapeutic conventional therapy [9]. In addition, these ACPs have been shown to quickly induce the death of cancer cells, hindering their ability to establish resistance [10].

Some ACPs are inherently antimicrobial peptides (AMPs) in nature [11] and have been classified according to secondary/tertiary formations: α-helical or β-sheet structures [12]. Owing to this dual property, cancer cells were predicted to have less resistance to cellular acquisition than that presented for chemotherapy [9]. In addition to those properties, AMPs that possess anticancer activity are characterized by having small amino acid sequences that range from 5 to 50 amino acids with a significantly low molecular weight.

In this study, we screened for a novel antimicrobial peptide with potential anti-cancer activity from the AUC Red Sea metagenomics data library generated during AUC/KAUST Red Sea microbiome project. The metagenomic samples primarily came from three locations in the Red Sea: Atlantis II Deep, Discovery Deep, and Kebrit Deep [13]. Scientists studying the chemical properties of the Atlantis II Deep and Discovery Deep brine pools observed the former was predominantly sulfur enriched, as opposed to the latter, which was carbon and nitrogen enriched. This difference in chemical composition resulted in unique microorganism diversity observed at the deepest soil layers in each site, with microorganisms specializing in metabolizing the compounds dominating the respective sites. A novel 37 residue antimicrobial peptide was identified from the microbiome of Atlantis II Deep brine pool. This peptide’s amino acid sequence was modified to increase its hydrophobicity and anticancer profile.

This 37-mer peptide was then tested in a dose-dependent cytotoxic response, on grade I, II-III/IV hepatocellular carcinoma cell lines, HepG2 and SNU449 respectively, ovarian cancer cell line, SKOV3, and HeLa cervical cancer cells. The anticancer properties of the peptide were further evaluated via the change in cell viability, morphology, and inhibition of cellular migration. The mode of cell death caused by the peptide was also investigated. Targeted selectivity of the peptide was additionally validated via screening of human erythrocytes. The peptide was also assessed for its antimicrobial activity on both gram-positive (*S. aureus*) and gram-negative (*E. coli*) bacterial strains, to verify its antimicrobial potential.

## Materials and Methods

### Metagenomics library screening and candidate peptide selection

We adapted a method previously proposed [11] for mining large datasets containing potential anticancer peptide sequences to search the AUC Metagenomics library. We compared a dataset of experimentally validated anticancer peptides [14,15,16,17,18] to a publicly available dataset of antimicrobial peptides and another of random peptides.

All possible oligopeptide frequencies were investigated in a size range from 1 to 30 amino acids. We also calculated the amino acid and dipeptide (2 amino acids) frequencies from the anticancer and antimicrobial datasets. We compared the frequencies of amino acids and dipeptides in the anticancer and antimicrobial peptide datasets to those of the peptides in the Metagenomics library (t-test, p<0.05, Bonferroni multiple testing correction). Only the peptides recognized as anticancer, according to their conformance to known amino acid frequencies, were selected. A sliding window of increasing size, starting from 5 amino acids, was used to generate the peptides from the Metagenomics library translated reads. The recognized peptides were scored according to their conformity with the mean amino acid and dipeptide frequencies from the experimentally proven anticancer peptide dataset, that is, they fall within the standard error from the mean. Additionally, we analyzed passing peptides for the presence of Hidden Markov Models (HMM’s) [19, 20] previously reported on experimentally verified anticancer peptides.

We confirmed our final predictions using the online tool developed by Tyagi et al [11] (http://crdd.osdd.net/raghava/anticp/).

#### Filtering a single candidate for modeling

A single anticancer peptide was selected from the final shortlist of 59 potential anticancer peptides for modeling. The criteria used were cationicity, model prediction score, and size.

#### Peptide performance optimization

To increase the statistical performance of our peptide, when run against a dataset of experimentally validated and random peptides, we carried out a series of optimization steps. We ran the peptide sequence as a FASTA file on the AntiCP web server for anticancer peptide prediction. We chose “model 2” analyses, where the peptide sequence query was compared to a set of experimentally validated anticancer peptides and random peptides. Amino acid modifications were serially introduced to maximize the prediction score. The modifications selected were those that occurred outside of the HMM alignment region. Serial modifications were concluded when the prediction model returned a score high enough to differentiate the query and identify it as an anticancer peptide.

#### Sequence alignments

We carried out two sequence alignments: DELTA-BLASTp (NCBI) and HMMER (EBI). The peptide sequence was queried in FASTA format on the DELTA-BLASTp platform. The *Homo sapiens* database was examined for all non-redundant proteins with a filter on the low complexity regions; all other parameters were set to their default values. We conducted as many iteration searches as needed until convergence, were no more new reference sequences aligned with our peptide. Any unknown or unnamed sequences were removed from each iteration.

We also used our peptide sequence to conduct an HMMER search using default parameters over the *Homo sapiens* reference genome. We iterated the search until convergence.

#### Predictive Modelling

We ran our peptide through I-TASSER, a web server for protein structure and function prediction [21]. Query submissions of the peptide sequence were done in FASTA format. Output files were extracted for further modeling and analysis. Modeling and visualization software Chimera, developed by UCSF, was utilized to produce all modeling and visualization figures and descriptions of the peptide structure [22]. We also utilized the PlifePred webserver to predict the estimated blood-borne half-life of our generated peptide sequence [23].

#### Peptide synthesis and preparation

Lyophilized powder of the peptide was synthesized by GL Biochem LTD, Shanghai, China, at 98% purity at a concentration of 1 mg per vial, stored in -20°C. Fresh stock of the peptide was prepared by dissolving the 1mg of the peptide in 1 ml of sterile deionized water (1 mg/ml).

### Cell culture

The following cell lines were used in this study: SNU449, HepG2, SKOV3, HeLa cells 1BR-hTERT. SNU449 cells were received as a gift from Dr. Mehmet Ozturk at the Department of Molecular Biology and Genetics, Bilkent University, Turkey. HepG2 cells were previously purchased from Vacsera, Egypt. SKOV3 cells were provided by Dr. Anwar Abdel Nasser, American University in Cairo. HeLa cervical adenocarcinoma cells and Immortalized human fibroblast cell line 1BR-hTERT cells were kindly provided by Dr. Andreas Kakarougkas, American University in Cairo, Egypt. SNU449 and SKOV3 cells were maintained in complete RPMI-1641 media (Lonza) supplemented with 10% heat-inactivated fetal bovine serum (FBS, Thermo Fisher Scientific) and 5% Penicillin-Streptomycin (Thermo Fisher Scientific). HepG2, HeLa, and 1BR-hTERT cells were kept in DMEM-HG (Lonza) completed with 10% heat-inactivated FBS and 5% Penicillin-Streptomycin. Cells were cultured in an incubator set at 37°C with 5% CO2.

#### SNU449 doubling time

Three thousand cells/ml, in fresh complete RPMI, growth rate was calculated at 6h, 24h, 48h, and 72h. Cells were trypsinized (Trypsin EDTA, 0.25%, phenol red, Lonza) for 4-7 minutes in a humidified incubator set at 37°C with 5% CO2. Trypsin was deactivated with fresh complete RPMI (cRPMI). Cells were collected after the addition of cRPMI to deactivate trypsin, via centrifugation at 500 Relative Centrifugal Force (RCF), 4°C for 5 minutes. After resuspension in cRPMI, a hemocytometer was used for cell counting using Trypan blue (Thermo Fisher Scientific) staining. 20µl of cell suspension was added to an equal volume of Trypan blue and loaded into the hemocytometer chambers. The cellular count was performed using the formula:

##### Theorem 1

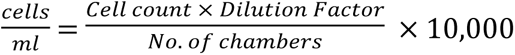

Cell count was calculated for each time point, and plotted on a growth curve using the following equation (ATCC®):

##### Theorem 2

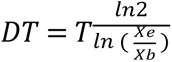

*DT* is the doubling time.

*T* is the total time during which exponential growth occurs.

*Xb* is the cell number at the beginning of exponential growth.

*Xe* is the cell number at the end of the exponential growth.

#### Cell cytotoxicity assay

SNU449, HepG2, SKOV3, HeLa, and 1Br-hTERT cells were seeded at 5×10^3^ cells/well in 96-well plate (Corning, USA), in fresh complete media (RPMI/DMEM, 10% FBS and 5% Pen-Strep). All cell lines were treated with peptide concentration gradient prepared in fresh complete media by serial dilution, for an exposure period of 24 h. Peptide concentration gradient for SNU449 cells was constructed for 24 h and 48 h time points and compared to a concentration gradient of cisplatin at 48 h.

Media with peptide was discarded and replaced with MTT reagent. Stock MTT reagent (5 mg/ml in PBS) (Serva, Germany) was diluted in complete media to reach a final concentration of 0.5 mg/ml. A diluted reagent was added to all conditions. Cells were incubated in the dark for 2-3 hours in a 37°C, 5% CO2 incubator, then media/substrate was discarded and replaced with DMSO (Sigma-Aldrich, USA), added to dissolve formed purple formazan crystals for 15 minutes in the incubator in complete darkness. Absorbance values were measured at 490 nm and 570 nm, with a BMG Labtech Spectrostar Nano plate reader. Blank corrected absorbance measurements reflected the cell viability percentage, with normalized untreated cell absorbance representing 100% viability.

#### Scratch wound healing assay

Scratch wound healing assay was performed to test the effect of the 37-mer peptide treatment on SNU449 and SKOV3 cell migration ability. SNU449 cells were seeded in duplicate at a density of 2×10^5^ cells/well in 12 well plates. SKOV3 cells were plated in 24-well plates (Corning, USA), at 5 ×10^4^ cells/well-fed with complete media. Two perpendicular lines were performed using sterile 20 μl pipette tips at 85-90% confluent monolayer cells. Following scratches, cells were gently washed twice with 1X PBS, to remove cell debris. Complete RPMI/DMEM media was added to the untreated control, while complete media was supplemented with IC50 of the peptide and added to the treated wells and incubated for 24 h. SNU449 cells were additionally assessed at 48 h treatment with the peptide due to its high calculated doubling time. Images of the scratch were taken at time points 0h, 24h, and 48h, and recorded with an Olympus IX70 inverted microscope. Wound closures were analyzed on ImageJ. Percentage of wound closure was calculated using the following equation:

##### Theorem 3

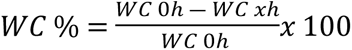

**WC** % = percentage wound closure (Cellular Migration).

**WC 0 h** = percentage wound closure at zero hours, once the scratch is performed.

**WC xh** = percentage wound closure at a specified time point.

#### Reverse Transcription Polymerase Chain Reaction (RT-PCR)

Gene expression for peptide-treated SNU449 and SKOV3 cells was analyzed using semi-quantitative RT-PCR for the following genes: KI67, B-catenin, Survivin, Bax, N-cadherin, E-cadherin, Vimentin, ATG5, ATG6, and ATG7. Extracted RNA (0.5 ug) was reverse transcribed using RevertAid First Strand cDNA Synthesis Kit (Thermo Scientific, USA), according to the manufacturer’s protocol. The PCR reaction was performed utilizing 1 µL cDNA template in Taq DNA Polymerase (Thermo Fisher Scientific) reactions. PCR conditions were standardized for tested genes: 94°C for 3 min, followed by cycles of 94°C for 30 s, annealing temperature for 30 s, 72°C for 45 s, with a final extension carried out at 72°C for 7 min. Additionally, each cycle number and annealing temperature for each gene was separately adjusted (Supplementary Table S1) based on optimization conditions. PCR products were run on 2% agarose gel and visualized in a Gel Doc EZ System (Bio-Rad, USA). GAPDH served as an endogenous control.

#### Annexin V apoptosis assay

To further study the peptide cytotoxic effect on the SNU449 cell line, Alexa Fluor 488 Annexin V/ Dead Cell kit (Thermo Fisher Scientific) was used to identify cellular death mode. Annexin and PI buffers were prepared according to the manufacturer’s instructions. Annexin V labeled with a fluorophore, Alexa Fluor® 488 dye, which is a human anticoagulant, has a high binding affinity for phosphatidylserine (PS). PS is exposed on the outer leaflet of the apoptotic cell plasma membrane while presented in the inner membrane of a viable cell. Necrotic cells are dyed with red fluorescence propidium iodide (PI) nucleic acid binding dye. PI can’t bind to apoptotic or viable cells.

SNU449 cells were plated at a density of 3×10^5^ per well in 6-well plates. After 24 hours, media was replaced by peptide’s IC50 complete media. At 24 h incubation, cells were collected via trypsinization. Cells were centrifuged for 5 minutes at 500 RCF, 4°C. Cells were resuspended in fresh media. The cellular suspension was mixed with Trypan blue counted via a hemocytometer. The suspension was centrifuged to an equivalent of 0.1×10^7^ cells/ml and washed with PBS. Cells were resuspended in 1x Annexin binding buffer and incubated for 15 min in complete darkness at room temperature. We visualized the cells on a slide under an Inverted microscope (Olympus IX70, USA), using the appropriate FITC filters.

#### Hemolysis assay

Blood samples were collected from healthy volunteers after acquiring an Institutional Review Board (IRB) approval and volunteers signed consent, anonymously. 2 ml of fresh blood was collected for every experiment by a physician in a clinical setting. Human erythrocytes were collected via centrifugation at 1000 g for 5 minutes and washed twice by PBS. Anticoagulant EDTA (Ethylenediaminetetraacetic acid) was used to keep erythrocytes dispersed. Following complete removal of serum, red blood cells were suspended in 2% PBS solution, where 50 µL of 2% erythrocytes were added to each well of a 96-well plate. Two peptide concentrations were used to test peptide hemolytic activity: IC50 and IC25. 50 µL of each concentration was dissolved in PBS and added to the 2% erythrocytes in the 96-well plate, to reach a final concentration of 1% human erythrocytes in each well. PBS was used as a negative control for hemolysis, while deionized water was used as a positive control. Plates were incubated at 37°C for 1 hour. Following incubation, plates were centrifuged for 10 min at 3000 g in a plate centrifuge, and the supernatant was transferred to a clean 96-well plate (flat bottom). Release of hemoglobin was detected via measuring the absorbance of the supernatant at 570 nm. Erythrocytes in PBS and deionized water were used as control of 0% and 100% hemolysis, respectively. Experiments were performed four times in triplicates.

The equation used to calculate Hemolysis percentage: [24]

##### Theorem 4

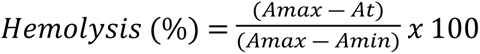

A_max_, A_min_, and A_t_ represented the absorbance values for untreated RBCs, completely hemolyzed RBCs, and tested RBCs.

#### Antimicrobial assay

The antibacterial activity of the peptide was tested on two bacterial strains: *Escherichia coli* (gram-negative strain) and *Staphylococcus aureus* (gram-positive strain). The overnight culture was prepared in nutrient broth from bacterial glycerol stock and kept in a shaking incubator at 37°C. 160 µl of the overnight culture was added to 8 ml of fresh media. Optical density (OD) was recorded at different time points, 0, 1, 2, and 3 hours, until the reading reached 0.1. At OD = 0.1, bacteria culture was in the log phase. Serial dilutions were prepared from culture, plated, and incubated overnight. Colonies were counted for 6 dilutions of each bacterial strain for a total of 106 CFU. Colony-forming units (CFU) were calculated at OD=0.1, with no dilutions. 50 µl of overnight culture was loaded in 96 well plates, in quadruplets, with 106 CFU in each well. Two peptide concentrations were prepared to be tested: 118.7 µM and 161.6 µM, the median and highest concentrations tested on SNU449. A broad-spectrum antibiotic (Cefoxitin®) was used as a positive control, with a final concentration of 30 mg/ml used in testing. Bacterial treated cultures were left overnight in a 37°C incubator. 20 µl from each condition was used to perform serial dilutions with clean broth and then plated on an agar plate for colony counting. Plates were incubated overnight. Absorbance was read at 600 nm.

Colony-forming unit equation:

##### Theorem 5

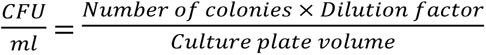

#### Statistical analysis

All data generated was presented as the mean ± standard deviation of three independent experiments, with at least three replicas, and was analyzed using GraphPad Prism 7 software. Multiple comparison analyses were performed using either two-way or one-way ANOVA (analysis of variance) followed by Dunnett’s post hoc multiple comparisons test. Pairwise analysis was performed using multiple t-tests analysis with multiple comparisons corrected by post hoc correction with the Holm-Sidak method. Dose-response curves and IC50 values were generated in GraphPad Prism 7 using the equation: Log inhibitors vs. Normalized Response-Variable slope *P* values less than 0.05 were considered significant (\**P*-value < 0.05, \*\**P*-value < 0.01, \*\*\**P*-value < 0.001, \*\*\*\**P*-value < 0.0001). R2 values represented the goodness of fit of the non-linear regression model utilized in the cell viability curves. Hill-slope coefficients were also included as an indicator of cooperativity of ligand-receptor binding, the affinity of the drug to bind to cellular receptors.

## Results

### In Silico prediction

#### Most promising peptide candidates contained homeodomain

A total of 59 candidate anticancer peptides were produced from the support vector machine (SVM) search of the AUC metagenomics library. The most promising peptide selected, TKEQKEQIAKATGLTTKQVRNWYVQLNASIKVMLTSI **(Table 1 “Original” column)** contained a homeodomain HMM profile (PFAM: PF00046.24), suggesting a similar role to homeoproteins. This 37-mer peptide was derived from Atlantis II Deep brine pool sub-seafloor sediment section 6 (available on the NCBI Sequence Read Archive, SRA). The peptide was modified by amino acid substitutions p.V32C, p.M33C, and p.S36C, outside the homeodomain region, for a final sequence of TKEQKEQIAKATGLTTKQVRNWYVQLNASIKVCMCSC. The modifications were introduced to increase the peptide’s SVM model performance scores and cell permeability **(Table 1 “modified” column)**.

**Table 1.** Chemical properties of the modified peptides.

|  | **Original** | **Modified** |
| --- | --- | --- |
| **Peptide length** | 37 | 37 |
| **HMMid** | Homeobox | Homeobox |
| **HMM accession** | PF 00046.24 | PF 00046.24 |
| **HMM start** | 1 | 1 |
| **HMM end** | 31 | 31 |
| **ACP/NON-ACP score** | -0.98 | 0.28 |
| **Hydrophobicity** | -0.18 | -0.2 |
| **Hydropathicity** | -0.39 | -0.34 |
| **Hydrophilicity** | 0 | -0.01 |
| **Charge** | 4 | 4 |
| **Mol. Weight** | 4334.71 | 4213.53 |

We ran a DELTA-BLASTp alignment that used an iterative position-specific scoring matrix (PSSM) using the NCBI Conserved Domain Database (CDD) as a reference. Upon convergence, we observed that the most significant hits aligning with the peptide were those with leukemogenic homolog protein (e-value = 2e-7, total score = 45, identity= 45%, cover = 64%) and MEIS2 (e-value = 3e-7, total score = 45, identity= 45%, cover = 64%). Congruent to the BLASTp alignments, we also ran an HMM-based alignment using the EBI’s HMMER web tool. Upon reaching convergence with the peptide query, we extracted a list of 65 significant alignments (E-value range from 2e-11 to 4.9e-4). The alignments occurred within the homeodomain structure, namely the homeobox_KN, PBC, and SIX1_SD architectures. The alignment domains indicate similarities with certain protein families such as the MEIS, PBX, and SIX (**Fig 1**).

**Fig 1.**
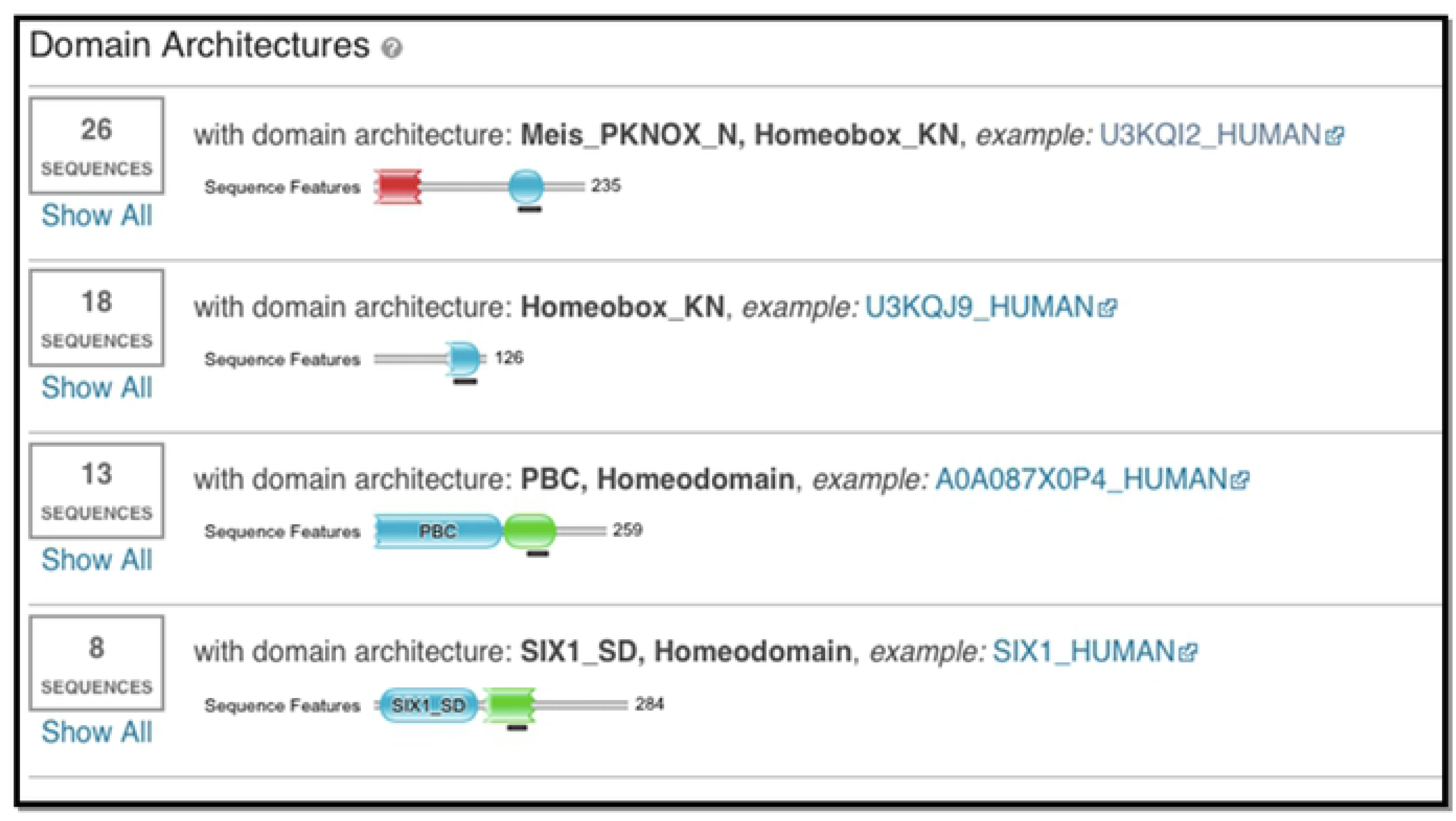
Domain architectures in which the HMMER alignments occurred against the peptide. The alignments occurred within the Meis_PKNOX_N, Hommeobox_KN, PBC, and SIX1_SD domain architectures.

#### Peptide contained helix coil structure similar to homeoprotein transcription factors

The sequence of the 37-mer peptide was input in the I-TASSER structure prediction software. We observed that the peptide contained a helix loop helix structure (**Fig 2A**); the helices formed a 79-degree angle between the axis of the C-terminal and N-terminal helix (**Fig 2B**). This architecture was consistent with the homeodomain consensus (PFAM).

**Fig 2.**
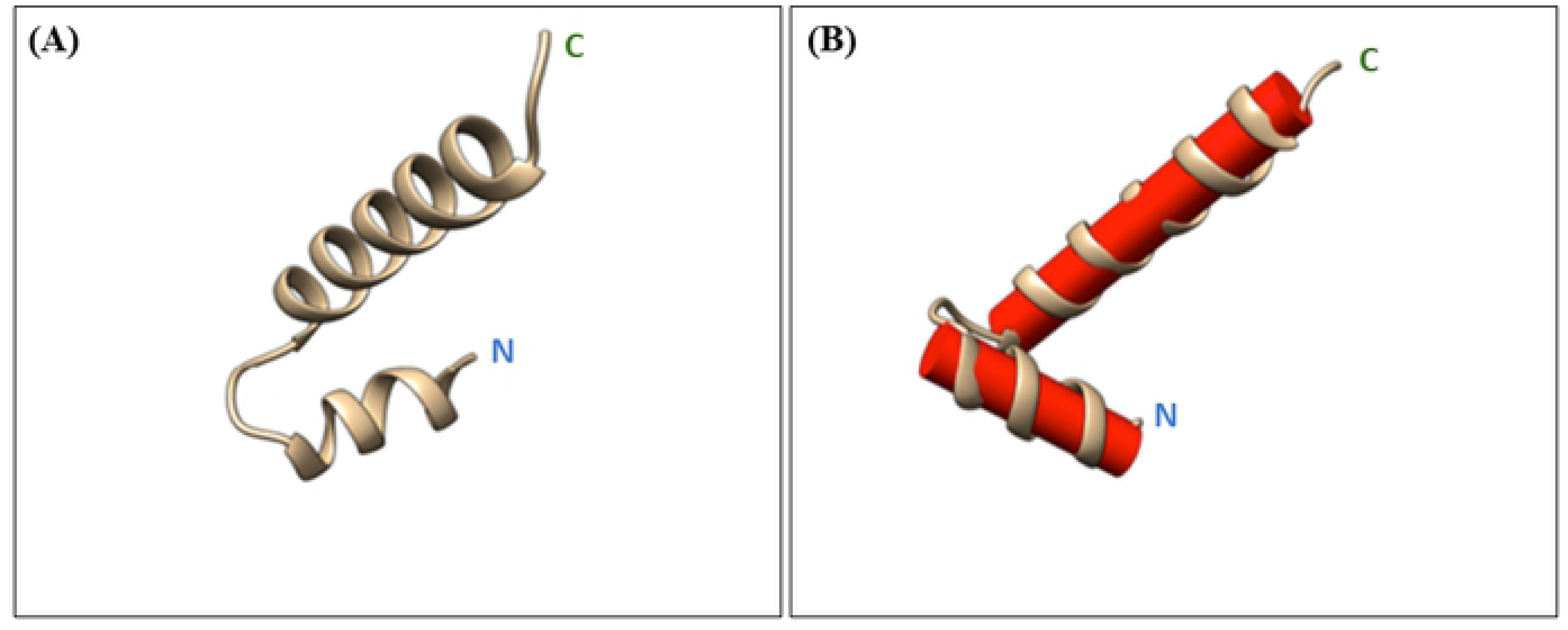
Peptide structure prediction results from I-TASSER. **(A)** Shows the secondary structure of peptide as predicted by I-TASSER. **(B)** Shows the angle formed between the axis of the C terminal recognition helix and the N-terminal helix which is about 79 degrees.

The DNA-bound structure of the extended PBX homeodomain and MEIS2 were used as templates to predict the structure of the peptide. Furthermore, we noticed that the protein showing the highest structural similarity to our peptide was the homeoprotein transcription factor PAX6 (TM-Score = 0.815, RMSD = 1.39, coverage = 1.000). Gene Ontology (GO) terms were generated for functional prediction by COACH analysis within the I-TASSER suite. We observed GO terms relating to sequence-specific DNA binding, non-covalent protein and protein complex binding, regulation of DNA transcription, and nuclear localization **(Table 2)**. Subsequent sequence analysis of modified peptide predicted half-life expectancy in the blood was 1325.81 seconds; equivalent to 22.1 min. Original peptide had a predicted blood level half-life of 1567.81 sec (26.1 min).

**Table 2:** GO-term annotations and functions predicted by I-TASSER for 37-mer peptide.

| GO-term | Function | Score |
| --- | --- | --- |

|  |  |  |
| --- | --- | --- |
| GO:0003700 | Sequence-specific DNA-binding transcription factor activity | 0.7 |
| GO:0043565 | Sequence-specific DNA-binding | 0.54 |
| GO:0005515 | Protein binding | 0.54 |
| GO:0006355 | Regulation of transcription | 0.7 |
| GO:0005634 | Nuclear localization | 0.33 |

### Cell culture analysis

#### MTT assay

MTT assay experimented peptide (37-mer peptide) dose-dependent cytotoxicity of SNU449, HepG2, SKOV3, HeLa, and 1BR-hTERT cells over serially diluted concentrations. A gradient of 71 µM to 162 µM was used to assess the peptide effect on SNU449 cells, while a lower series of concentrations ranging from 3.8 µM to 121.5 µM was used to test peptide treatment of HepG2, HeLa, and 1BR-hTERT cells for 24 h. A broader range of concentrations was tested with SKOV3, from 0.024 µM to 120 µM. Cellular viability decreased significantly, for all tested cell lines, with each increase in the peptide concentration, when compared to the untreated cells (**Fig 3**).

**Fig 3.**
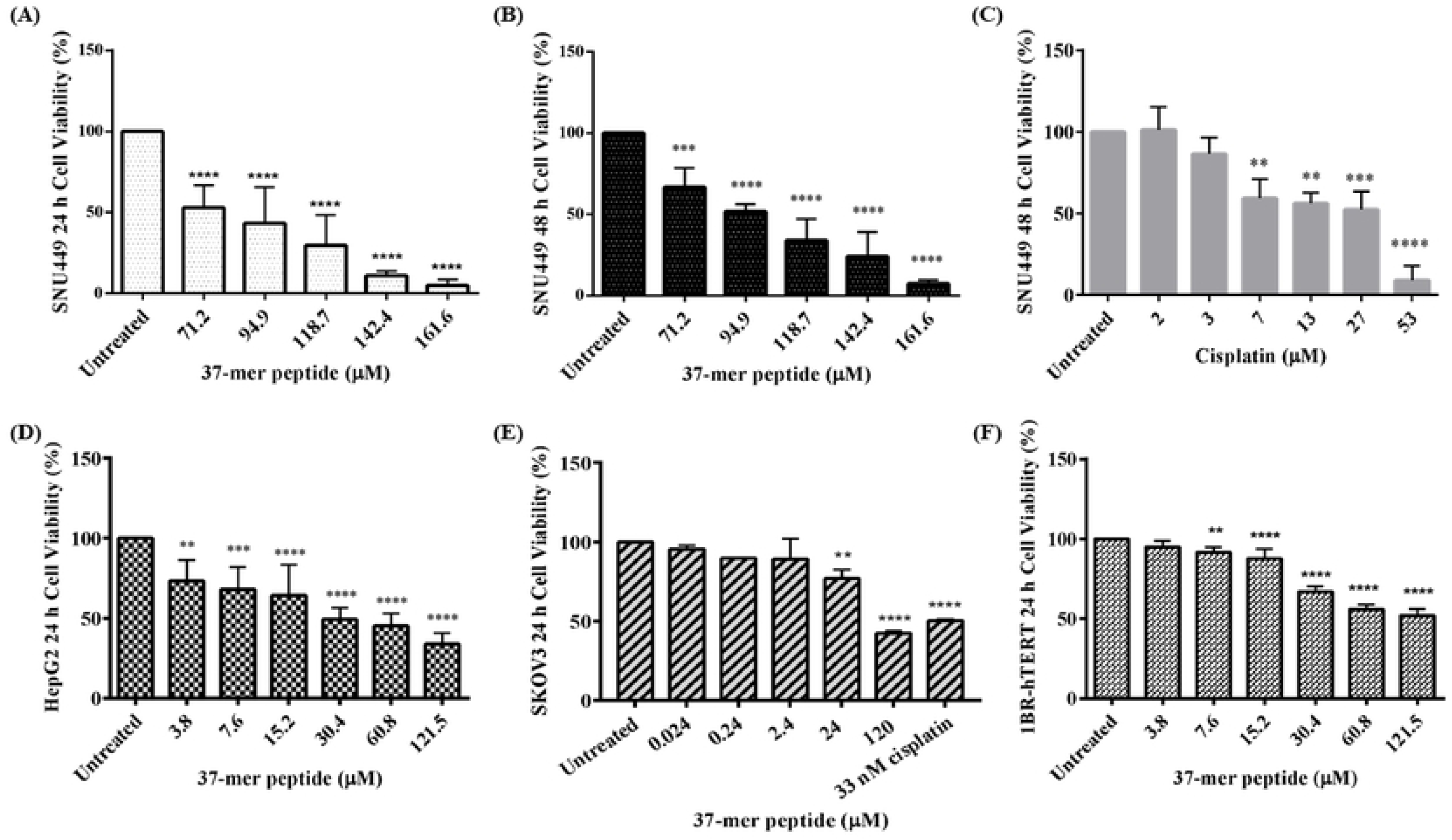
37-mer peptide dose-dependent cytotoxicity. **(A)** SNU449 displayed a significant decrease in cell viability (approx. 50 %) upon treatment with peptide (71.2 µM) at 24h. **(B)** Treatment of SNU449 for 48 h with peptide showed a similar trend to 24 h treatment. Cells also experienced significant cell death (approx. 60% viability) at 71.2 µM exposure. **(C)** SNU449 cells were screened with 48 h treatment of Cisplatin. 7 µM of cisplatin treated SNU449 showed similar cell viability to peptide treatment (71.2 µM). **(D)** HepG2 peptide treatment at 24 h displayed similar viability (approx. 50%) at a lower concentration (30.4 µM) compared to more advanced hepatocellular carcinoma SNU449 cells (71.2 µM). **(E)** SKOV3 treated with peptide for 24 h showed a decrease in cell viability (approx. 80%) at 24 µM. Treatment with 33 nM of Cisplatin reduced cell viability by approx. 50%. SKOV3 displayed greater sensitivity to the peptide, as cytotoxicity started to decrease from 0.24 µM of treatment. **(F)** 1BR-hTERT cells were treated with the peptide for 24 h. Cells showed similar viability (approx. 50%) at a lower treatment concentration (60.8 µM) compared to SNU449 cells (71.2 µM) (** P < 0.01, *** P < 0.001, **** P < 0.0001, n=3).

At 162 µM of peptide treatment, SNU449 cells showed an average of 4.8% viability, while 71 µM treatment showed an average of 52.8% viability at 24 h. Peptide-treated HepG2 cells showed a statistically significant reduction in viability at the lowest tested concentration, 3.8 µM. SKOV3 displayed greater sensitivity to the peptide, as cytotoxicity started to decrease from 0.24 µM of treatment. Upon treatment of HeLa cells with 37-mer peptide, we were unable to observe a consistent cytotoxic peptide response **(S1 Fig.**). 1BR-hTERT cells exhibited a statistically significant drop in cell viability starting at 15.2 µM (P<0.05), at the 24 h time point.

Cisplatin, utilized as a positive control, was evaluated on SNU449 cells specifically, with a concentration gradient ranging from 2 µM to 53 µM for 48 h treatment. High cisplatin concentrations, 53 µM, and 27 µM caused a significant reduction in SNU449 cells’ viability, coming at 8.8% and 52.3%, respectively. Peptide treatment for 48 h at 162 µM peptide concentrations caused an average of 7.2% viability, while 71 µM of the peptide caused an average of 66.7% viability.

#### The half-maximal inhibitory concentration (IC50)

Using nonlinear regression analysis, results obtained from MTT assay were directed to calculate peptide’s relative and absolute IC50, R2, and Hill-slope coefficients for SNU499, HepG2, SKOV3, and 1BR-hTERT cells for 24 h time point **(Table 3)**. Dose-response curves for all peptide concentrations were plotted for 24 h and 48 h (SNU449) to calculate IC50 for peptide treatment (**Fig 4**). HeLa data was similarly displayed **(S2 Table)**.

**Fig 4.**
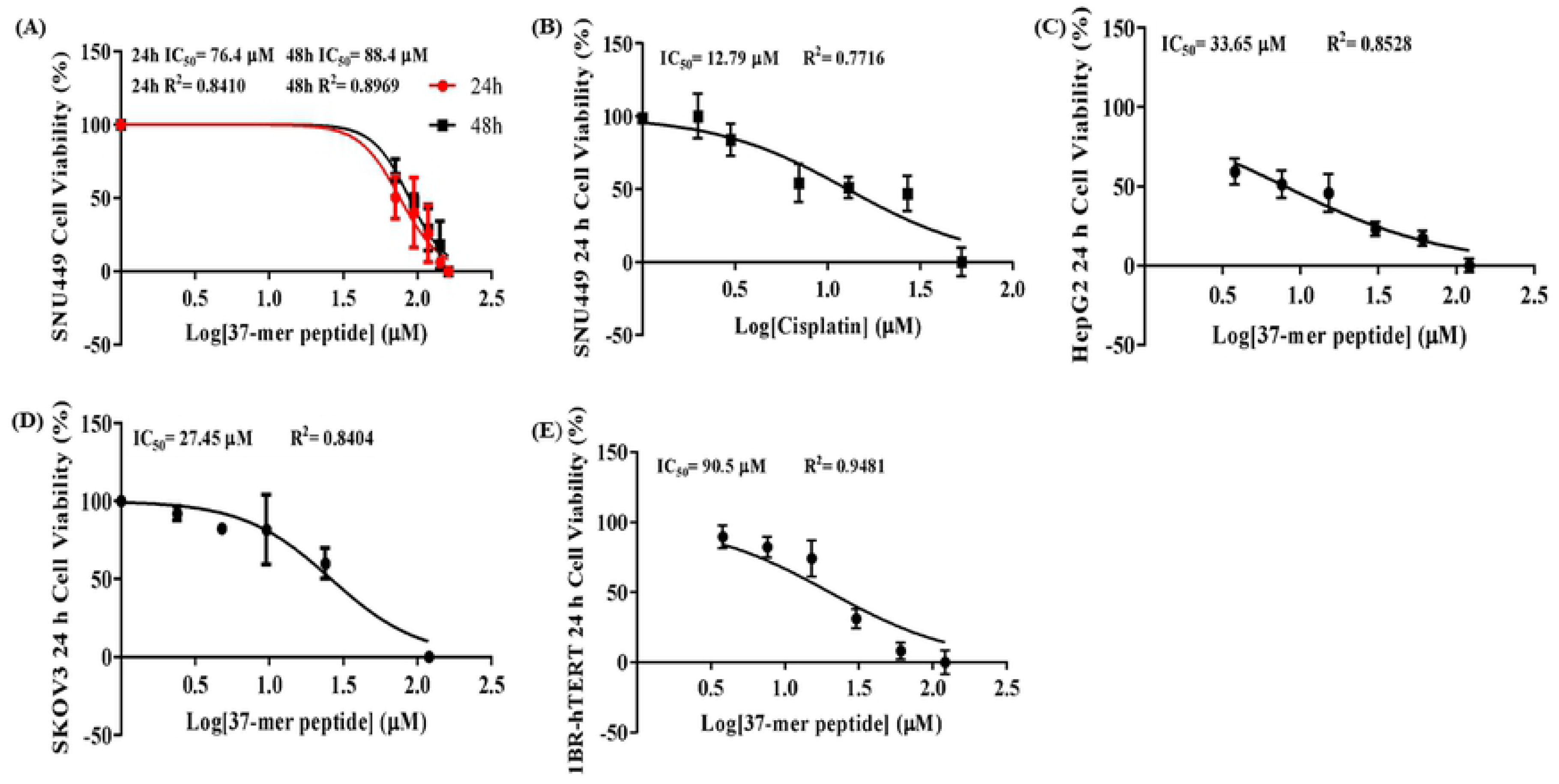
Dose-response of SNU449, HepG2, SKOV3, and 1BR-hTERT upon 37-mer peptide treatment. Cell response of each specific cell type was normalized to a scale running from 0 to 100. **(A)** Peptide treatment response of SNU449. IC50 value was calculated as 76.4 ± 0.6015 µM (R2= 0.8410) from 24 h treatment, and 88.4 ± 0.9148 µM (R2= 0.8969), from 48 h exposure, respectively. **(B)** Cisplatin exposure of SNU449 at 48 h produced a response curve from which the IC50 value was calculated as 12.79 ± 2.055 µM (R2= 0.7716). **(C)** HepG2 peptide 24 h treatment response curve produced an IC50 of 33.65 ± 1.09 µM (R2 = 0.8528). **(D)** SKOV3 response from 24 h peptide treatment produced an IC50 value of 27.45 ± 1.5085 µM (R2= 0.8404). **(E)** 1BR-hTERT 24 h dose-response curve generated an IC50 of 90.5 ± 0.521 µM (R2 = 0.9481).

**Table 3.** Summary of IC50, hill coefficient and R2 values obtained for peptide treatment for all cell lines.

| Drug | Cell line | Time Point (h) | Absolute IC<sub>50</sub> (μM) ± Std. Error | Relative IC<sub>50</sub> (μM) ± Std. Error | Hill Coefficient ± Std. Error | R<sup>2</sup> | Sy.x |
| --- | --- | --- | --- | --- | --- | --- | --- |
| 37-mer peptide | SNU449 | 24 h | 79.4 | 76.4 ± 2.6015 | 3.195 ± 0.6 | 0.8410 | 14.59 |
| 37-mer peptide | SNU449 | 48 h | 93.1 | 88.4 ± 0.9148 | 3.56 ± 0.44 | 0.8969 | 11.24 |
| Cisplatin | SNU449 | 48 h | 13.7 | 12.79 ± 2.055 | 1.183 ± 0.1789 | 0.7716 | 16.36 |
| 37-mer peptide | HepG2 | 24 h | 35.9 | 33.65 ± 1.09 | 1.169 ± 0.17 | 0.8528 | 18.54 |
| 37-mer peptide | SKOV3 | 24 h | 28.4 | 27.45 ± 1.5085 | 2.67 ± 2.699 | 0.8404 | 14.57 |
| 37-mer peptide | 1BR-hTERT | 24 h | 90.8 | 90.5 ± 0.521 | 2.058 ± 0.1958 | 0.9481 | 9.03 |

The relative IC50 for SNU449 was 76.4 µM (321.7 µg/ml) for 24 h time with R2 value 0.8410, and 88.4 µM (372.7 µg/ml) for 48 h time with R2 of 0.8969. IC50 for HepG2 cells was calculated at about 33.7 µM (141.6 µg/ml) with an R2 of 0.8528. SKOV3 absolute IC50 was calculated at approximately 27.5 µM (115.2 µg/ml) with an accompanying R2 of 0.8404. The relative IC50 for 1BR-hTERT cells was calculated at about 90.5 µM (380.9 µg/ml) for an R2 of 0.948.

#### Cell morphology

SNU449 and HepG2 cells exhibited morphological changes with each peptide concentration increasing in a dose-dependent manner. The epithelial morphology of SNU449 cells transformed to compact round cell structures with vacuoles, upon treatment. At higher concentrations (142.4 µM) of the treatment peptide, multiple fields of SNU449 cells showed extensive membrane damage resulting in cellular rupturing and release of intracellular components into surrounding media (**Figs 5A** **and 5B).** The remaining cells displayed shrunk cytosolic cell formations from peptide treatment (**Figs 5A** **and 5B)**. At 71 µM, the lowest concentration, most cells appeared viable; however loss of cellular content was still observed. Cisplatin effect on cellular morphology was observed and compared to peptide effect on SNU449 at 48h (**Fig 5C**). Cisplatin-treated cells showed circular formations with more noticeable cellular membrane compared to peptide-treated cells. At the highest concentration, peptide treatment of HepG2 resulted in enlarged cells with irregular shaping and rough membranes (**Fig 5D**), in addition to the fragmentation formed once treated with 60.8 µM. SKOV3 cells also displayed the release of multiple large vacuoles that resembled some of the treated HepG2 cells at the highest concentration (**Fig 5E**). Treated 1BR-hTERT cells seemed to be shrunken and had a rough outer membrane when compared to the untreated cells (**Fig 5F**).

**Fig 5.**
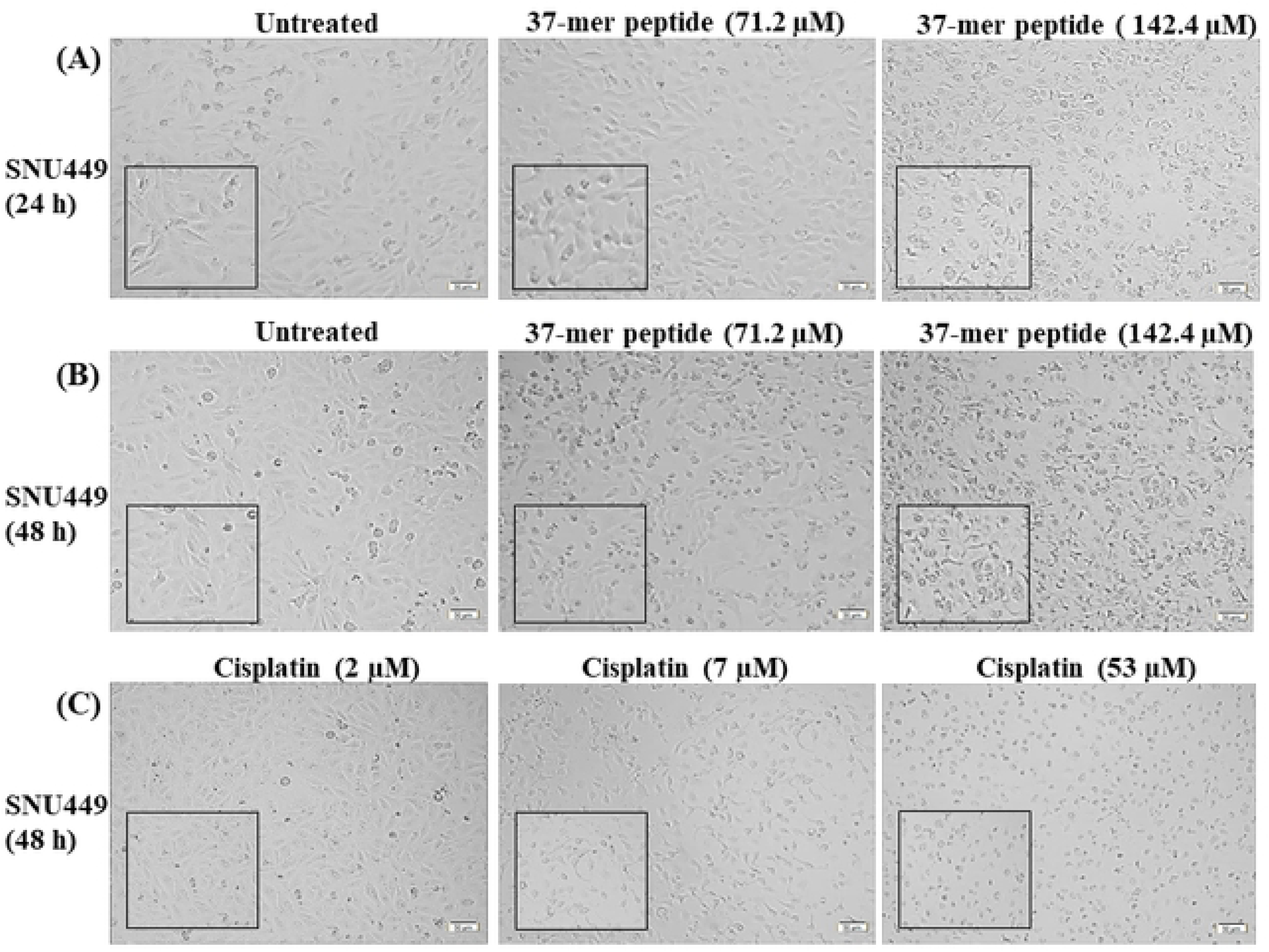
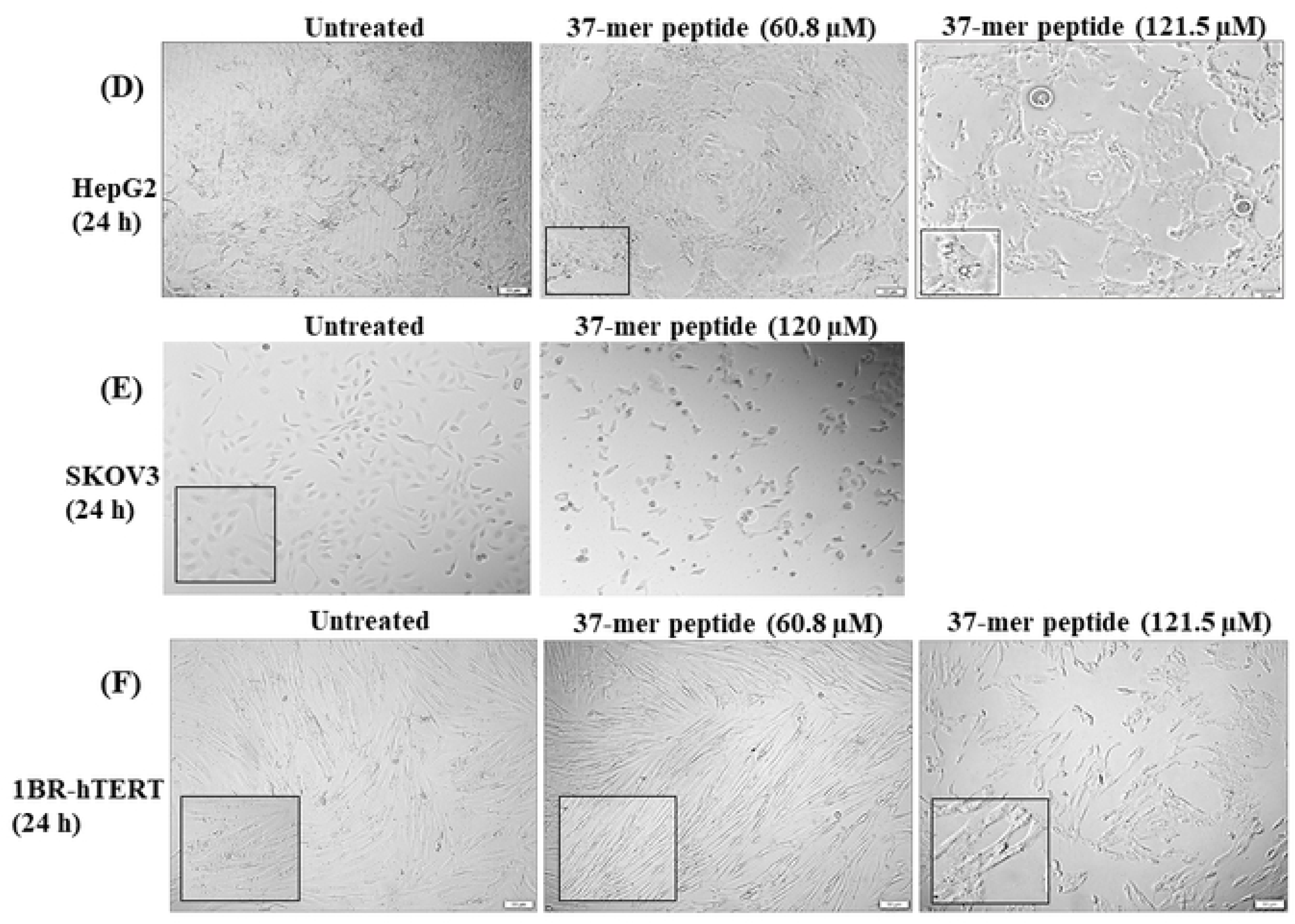
Peptide effect on cellular morphology of SNU 449, 1Br-hTERT, HepG2, and SKOV3 cell lines. **(A)** The lowest (71.2 µM) and the highest (142.4 µM) concentrations of peptide treatment on SNU449 cell line at 24 h. Untreated SNU449 cells are compact small spindle-shaped adherent epithelial cells. Treatment with 71.2 µM 37-mer peptide showed cells forming circular shapes and detachment of adherence surface. At higher peptide concentration, cells appeared to be surrounded by debris and form condensed cell detachments. **(B)**The lowest (71.2 µM) and the highest (142.4 µM) concentrations of the peptide on SNU449 cell line at 48 h treatment. Untreated cells in 48 h culture appeared similar to 24 h culture. Treatment with 71.2 µM of the peptide exhibited cells forming rounded sparse distribution. A higher concentration of treatment (142.4 µM) led to the aggregation of ruptured cells forming circular floating detachments. **(C)** Three concentration points in Cisplatin treatment gradient of SNU449 cells for 48 h: 2 µM, 7 µM, and 53 µM concentration. Induction of 2 µM cisplatin had minimal effect on SNU449 cell morphology compared to untreated cells. 7 µM Cisplatin treatment appeared to halt cell proliferation without changing cellular morphology. High concentration (53 µM) of cisplatin showed excessive cell rounding and reduced cell count. **(D)** The lowest (60.8 µM) and the highest (121.5 µM) concentrations of 37-mer peptide treatment on HepG2 cell line at 24 h. Untreated HepG2 cells formed a monolayer of an irregular epithelial-like shape. Treatment of HepG2 cells with 60.8 µM caused cells to become sparse, with fragmented growth, while not showing signs of cell rounding. With the highest concentration of treatment, the HepG2 monolayer appeared to be sparse and significantly more fragmented than untreated cells. Additionally, highlighted section included rounded and floating cells, surviving cells do not resemble original cell morphology. **(E)** SKOV3 cell images of untreated and peptide-treated cells (120 µM). SKOV3 cells formed smaller circular spindle square formations. Administration of 120 µM of peptide caused cells to change morphology and become sparse, circular, and condensed, characteristic of dead cells. The highlighted section shows rounded detached morphed cells with surrounding debris. **(F)** The lowest (60.8 µM) and the highest (121.5 µM) concentrations of peptide treatment on 1BR-hTERT cell line at 24 h. Untreated cells formed elongated, aligned spindle-shaped morphology. The treatment of 1BR-hTERT cells with 60.6 µM appeared to break up cells without causing visible cell death. High concentration (121.5 µM) exposure displayed fragmented cells while losing the spindle elongated morphology, with some cells forming circular shrink appearance.

#### Scratch-wound healing assay

A scratch wound healing assay was performed to test the effect of peptide treatment on SNU449, and SKOV3 cell migratory ability. We observed a significant effect on the capabilities of both cell lines to close peptide-treated wounds. The wound gap increased by 2% and 4% for SNU449 cells exposed to the peptide at 24 h and 48 h, respectively, compared to untreated cells, where the wound gap decreased by 15% and 43% for 24 h and 48 h, respectively (**Fig 6**). Thus, peptide treatment induced a statistically significant reduction in SNU449 migration for both time points. SKOV3 cells displayed wound closures of around 51.18% when untreated, while peptide treatment significantly reduced migration to account for 15.15% wound closure.

**Fig 6.**
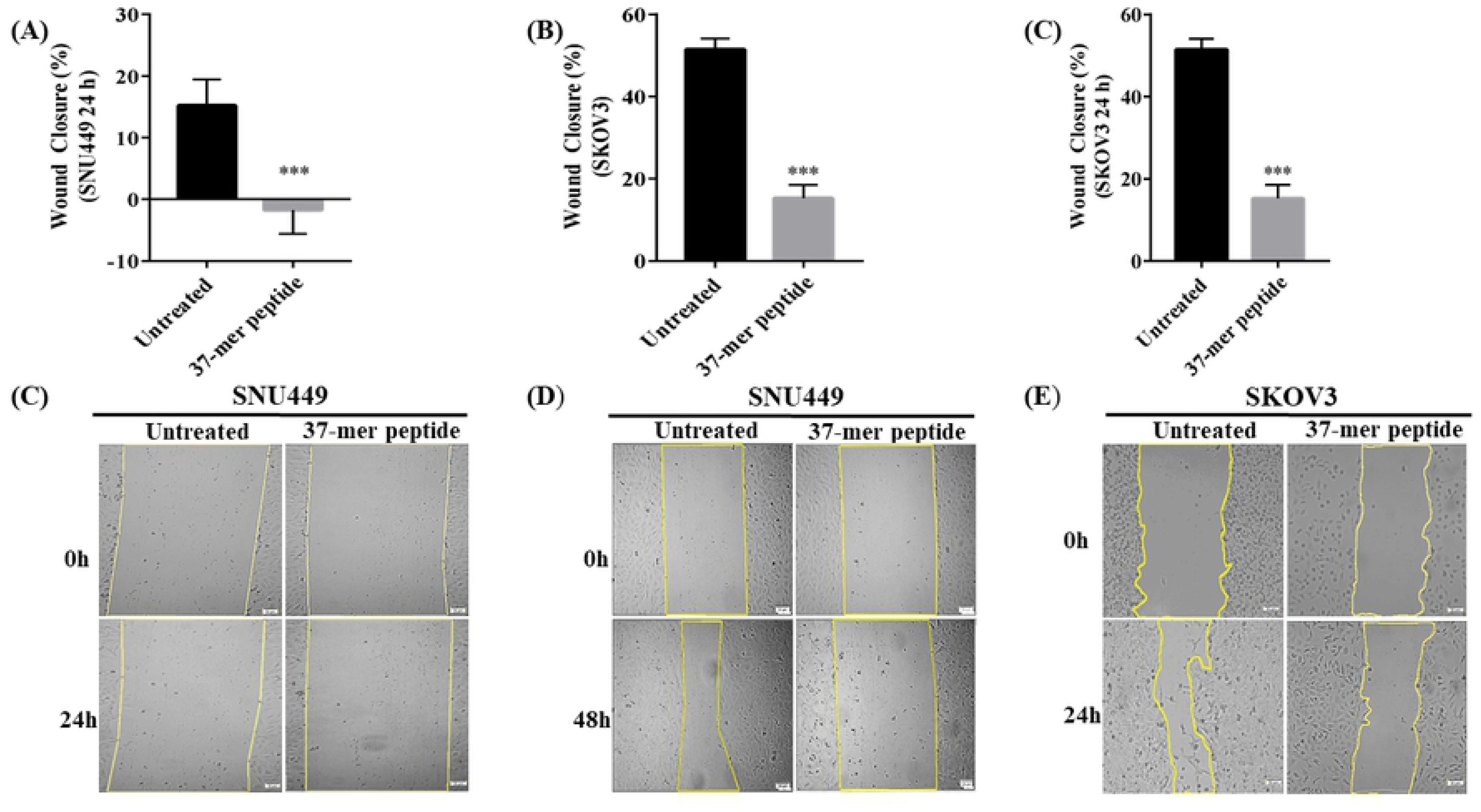
37-mer peptide treatment hindered SNU449 and SKOV3 cell migration. Highly confluent cells were subject to vertical scratches to observe cell migration effect when treated with IC50 of the peptide. **(A)** and **(B)** Percentage wound closure of SNU449 cells measured at 24 h and 48 h time points at the IC50 concentration respectively. Wound closure decreased by approximately 20% between untreated and treated (76.4 µM) cells, in addition, the closure also diminished by approximately 50% following 48 h exposure (88.4 µM). **(C)** Average wound closure of untreated SKOV3s and 37-mer peptide treated (27.5 µM) cells. Treated SKOV3 closure was significantly reduced, by up to 40%, compared to untreated cells (*** P < 0.001, **** P < 0.0001, n=3). **(D)** Representative wound closure images of SNU449 cells treated with peptide for 24 h. **(E)** Similarly, shows wound healing images of SNU449 cells treated for 48 h. **(F)** Wound closure of SKOV3 cells untreated and treated with peptide.

#### Gene expression

SNU449 and SKOV3 cells treated with the peptide for 24 h were profiled for specific gene expression levels associated with proliferation, epithelial to mesenchymal transition (EMT), apoptotic, autophagic, and survival-oriented genes. SNU449 cells were examined for KI67, B-catenin, Survivin, Bax, N-cadherin, E-cadherin, Vimentin, ATG5, ATG6, and ATG7. Cells did not show any significant effect on gene expression levels **(S2 Fig).** SKOV3 cells were profiled for KI67, B-catenin, Vimentin, ATG5, ATG6, and ATG7. Peptide treatment caused a significant reduction in KI67 and B-catenin expression while increasing Vimentin and ATG6 expression compared to untreated cells **(S3 Fig)**. However, ATG7 expression was inhibited in treated cells.

#### Annexin V apoptosis assay

Annexin V assay was utilized to assess SNU-449 cell line mode of death, upon peptide treatment at 24 h time point. Induction of the peptide caused an increase in the number of early apoptotic cells for SNU449 cells at a 24 h time point (**Fig 7**). The number of cells in early apoptosis was significantly higher in peptide-treated cells than in the untreated control. The high number of cells in early and late apoptosis suggested that most cells indeed underwent apoptosis or membrane rupturing. In treated cells, 51% of cells were in early apoptosis, 18% of cells were in late apoptotic/necrotic and 31% were live cells. With untreated control, 6% of the cells were in early apoptosis, 21% were in late apoptosis/necrosis and 72% of the cells were viable.

**Fig 7.**
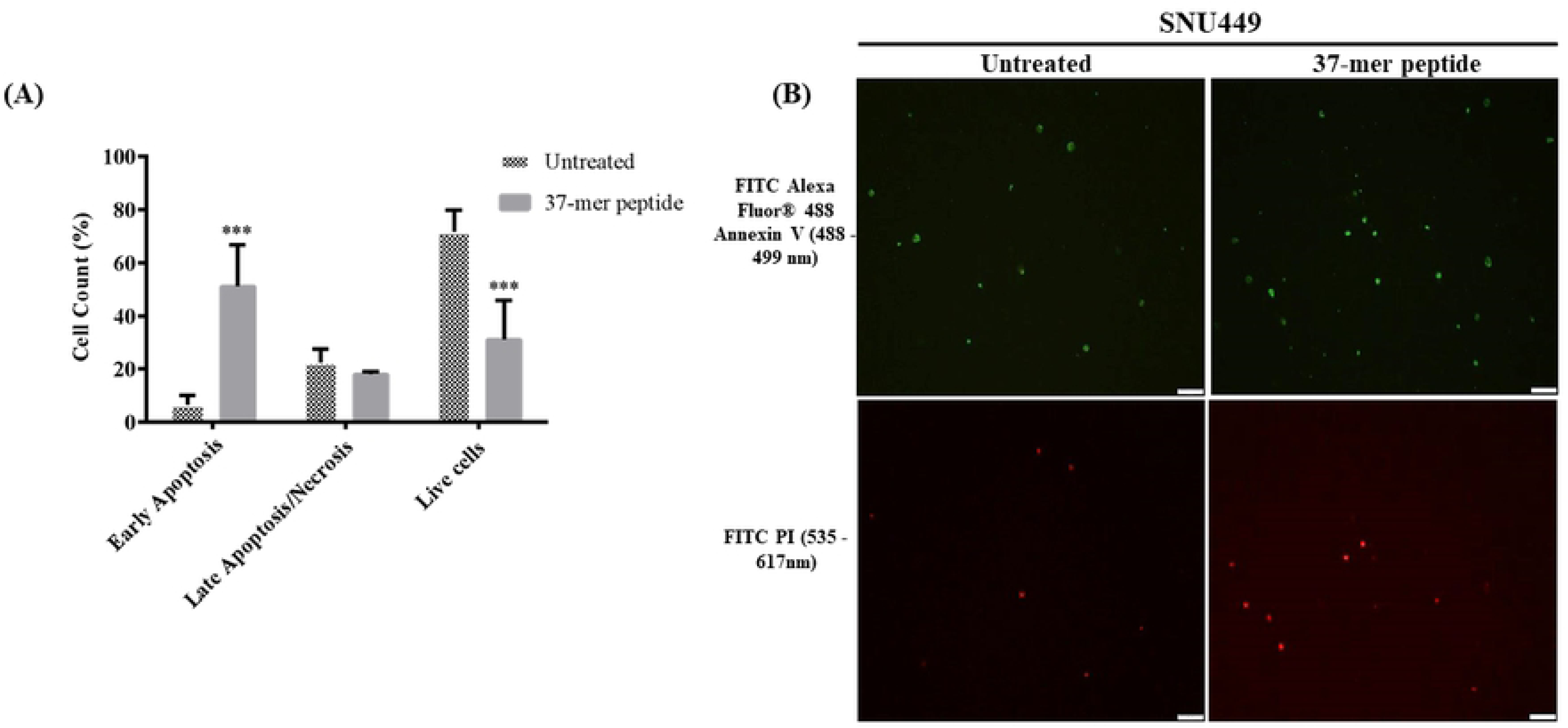
37-mer peptide IC50 treatment apoptotic effect on SNU449 at 24h. **(A)** Treated cells exhibited early-stage apoptotic death, no effect on late-stage apoptosis/necrosis, with a significantly lower live cell count (*** P < 0.001, n=3). **(B)** Representative images of Annexin V assay microscopy. Merged images showing overlapping green and red signals in untreated control and 37-mer peptide IC50 treated fields. Peptide treatment appeared to increase the rate of early apoptosis in SNU449 cells, compared to untreated cells. Additionally, PI staining was visually similar whether cells were treated or not.

#### Hemolysis Assay

Hemolytic activity of the 37-mer peptide was assessed by applying the IC50 (76.4 µM) and IC25 (53.8 µM), from peptide treatment of SNU449, on 2% human erythrocytes. Hemolytic assay measured the amount of hemoglobin liberated from lysed erythrocytes by spectrophotometric means. IC25 of the peptide caused 2.7% hemolysis to RBCs, while IC50 caused 5% hemolysis. Saline resulted in 2.7% hemolysis (**Fig 8**). Saline was used as a solvent to prepare peptide concentrations. The hemolytic effect of saline was normalized against the hemolytic effect of peptide, resulting in the negligible hemolytic activity of IC25 and IC50 treatment.

**Fig 8.**
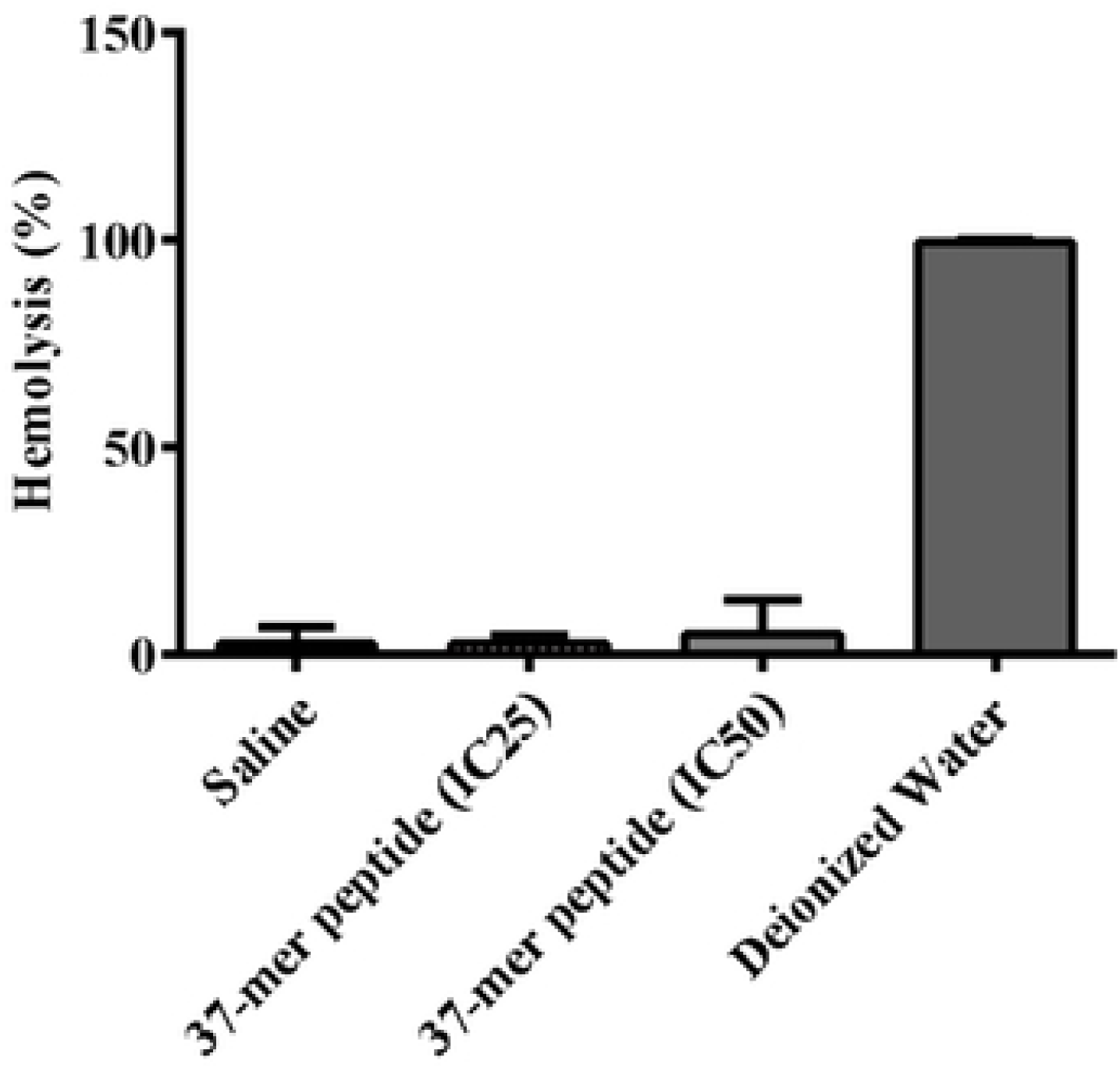
37-mer peptide treatment showed minimal hemolytic activity. The hemolytic activity of SNU449 IC25 (53.8 µM) and IC50 (76.4 µM) concentrations were determined via incubation of erythrocytes for 1 h. Peptide treatment did not show a statistically significant effect on erythrocytes, compared to saline. Deionized water represented standard positive control with 100% hemolysis. Saline represented negative control, n =4.

#### Antimicrobial activity assay

The antimicrobial assay was used to assess the antimicrobial activity of the peptide on gram-negative and gram-positive bacterial strains. The median and highest peptide concentrations tested on SNU449 cells (118.7 µM and 161.6 µM) were added to *S. aureus* and *E. coli* treatment, to examine the peptide’s broad-spectrum antimicrobial activity (**Fig 9**). The antimicrobial effect of these concentrations was tested by recording the absorbance of the turbidity of samples following treatment. Peptide concentrations 118.7 µM and 161.6 µM caused 44% and 49% viability in *S. aureus* and 56% and 42% viability in *E. coli*, respectively. A broad-spectrum antibiotic was used as a positive control for both microorganisms. Cefoxitin®, 30 mg/ml, caused 10% viability in *S. aureus* and 40% viability in *E. coli*. The peptide effect was also assessed as a function of the number of colonies formed in culture (CFU/ml). Following 24 h incubation samples were serially diluted and plated on agar plates. The peptide treatment with 118.7 µM and 161.6 µM produced a decrease in the number of colonies of both bacterial strains in agar plates. The fourth dilution gave the best visual counting field for colonies. *S. aureus* samples treated with 118.7 µM resulted in the survival of 52% of CFU/ml and treatment with 161.6 µM resulted in the survival of 22% of CFU/ml. The exposure of *E. coli* to 161.6 µM concentration of the peptide resulted in the survival of 20% of the CFU/ml, while 118.7 µM induction caused the survival of 5% CFU/ml. Overall the peptide showed statistically significant antimicrobial activity on both gram-negative and gram-positive bacterial strains.

**Fig 9.**
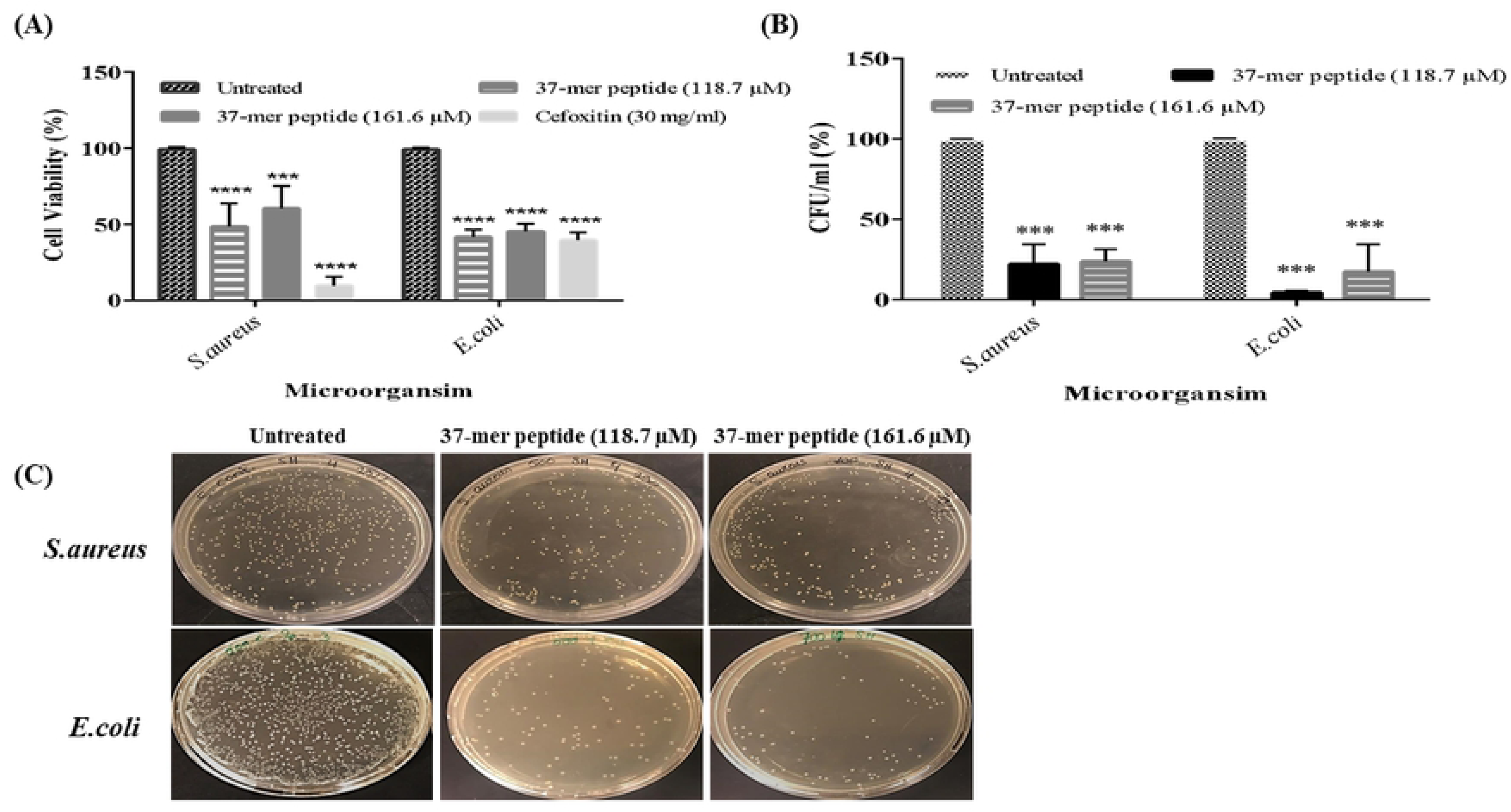
The antimicrobial activity of *S. aureus* and *E. coli* was evident with 37-mer peptide treatment. A statistical representation of peptide antimicrobial activity on *S. aureus* and *E. coli*. **(A)** Effect of 161.6 µM and 118.7 µM of peptide treatment on viable *S. aureus* and *E. coli* bacteria using microdilution assay. Cell viability was determined via absorbance measurements at 600 nm. Cefoxitin (30 mg/ml) was considered a positive control. Peptide treatment resulted in a significant loss of viability, approximately 50% or lower, in both gram-positive (*S. aureus)* and gram-negative (*E. coli)* bacterial strains. **(B)** The colony formation unit of *S. aureus* and *E. coli* cells was significantly debilitated (approximately 25%) by both concentrations. *S. aureus* specifically, showed an approximate 20% (118.7 µM) and 25% (161.6 µM) decrease in CFUs overnight exposure, respectively. *E. coli* CFU count displayed an approximate 5% and 15% respective decrease with medium and high treatment, respectively (*** P <0.001, ** P <0.01, n=3). **(C)** Representative images of agar plates showing viable colonies of *S. aureus* and *E. coli* after overnight culture (10^6^ dilution factor). A statistical representation of peptide antimicrobial activity on *S. aureus* and *E. coli*.

## Discussion

In this study, we developed an SVM workflow for the detection of anticancer peptides from the AUC Red Sea metagenomics library. Our SVM returned a list of potential anticancer peptides each with a model performance score. We selected the best candidate peptide out of the original list of 59 peptides, using cationicity and HMM domain as the main criteria of selection. Furthermore, serial modifications were introduced to the chosen peptide sequence outside of the homeodomain region, to enhance its model performance score as an ACP. Modifications did not significantly decrease predicted blood-borne half-life. We believe that our SVM model initially could not recognize any peptides as ACPs because there was very little variation between the ACP/Non-ACP groups, thus a larger sample size with more diversity was required to create a clear distinction. In other words, the model can discern ACPs from AMPs due to the distinct properties of ACPs; however, compared to all random peptides, a much larger data set would be required to train the SVM to recognize anticancer peptides in a pool of random peptides.

To gather more information about our peptide, we ran two sequence alignments. The first sequence alignment was conducted using NCBI’s DELTA-BLASTp alignment tool, the second was conducted using EBI’s HMMER alignment. We observed that our peptide consistently aligned with the homeodomain region of several homeoproteins from the MEIS, PBX, and SIX families. These observations all corroborate the previous finding that our peptide contains a homeodomain.

Secondary structure prediction using the I-TASSER suite indicated that the peptide contained two α-helical regions with architecture consistent with that of the DNA binding region of the homeodomain. The ligand-binding prediction apparatus of the I-TASSER suite predicted that our peptide would bind to DNA fragments. GO-term annotation pointed towards the potential implication of our peptide in pathways involving the direct sequence-specific binding of DNA and modulation of gene transcription.

Through MTT experiments we observed that our peptide was able to elicit a cytotoxic response in SNU449, HepG2, and SKOV3 cells. Treatment of HeLa cells appeared to cause numerous large vacuole formations, with partial cytotoxicity. Treatment of human 1BR-hTERT cells did induce a cytotoxic response; however, more was attenuated compared to SNU449 or HepG2 cells. The large peptide IC50 value calculated for 1BR-hTERT cells indicates that our peptide has the potential to be selective towards cancer cells while having a less pronounced effect on non-cancer cells. Hill-slope values were all above 1, indicative of a positive cooperative feedback loop of the binding of the peptide.

Cisplatin treatment, used as an anticancer control drug, was evaluated on SNU449 at 48 h and compared to the effect of the peptide. Cisplatin showed an increased cytotoxic effect on SNU449 than that of the peptide. IC50 of peptide treated SNU449 cells was calculated to be 88.4 µM, compared to 16.4 µM for cisplatin. Comparing 24 and 48 h peptide treatment of SNU449, higher peptide IC50 at 48 h compared to 24 h (76.4 µM) might suggest that peptide molecules were consumed by cells at 24 h and surviving cells were able to recover and proliferate. Thus, an increase in peptide concentration was required to provide an effective dose-response.

Upon treatment, peptide caused an irreversible morphological change to SNU449, HepG2, SKOV3, and 1BR-hTERT at 24 h time point. SNU449 cells were altered from an epithelial diffused cell structure to form more compacted circular cells releasing vacuoles resembling autophagosomes, with partial rupture and collapse upon the nucleus. With increasing peptide concentration, the damage in the cellular morphology was observed to be irreversible, with cells undergoing multiple death pathways. Vacuole formation was also visible in treated HepG2 cells. Major loss in the normal cellular topology was identified in treated cells, specifically more distinct in SKOV3 cells that became significantly more fragmented and deformed.

The primary two cell lines of interest, based on the peptide dose-dependent response, are SNU449 and SKOV3, where IC50 values of each cell line were calculated to be the highest and lowest, respectively, compared to other tested cells. Thus, to further characterize the peptide’s cytotoxic and debilitating effect on these two cell lines, cell migration was evaluated using a scratch wound-healing assay. Due to the drastic changes in cellular morphology and shrinkage observed in cells on the sides of the scratches, SNU449 cells lost their ability to migrate, thus resulting in the enlargement of the scratches formed, compared to untreated controls in both 24h and 48h exposure conditions. Additionally, SKOV3 wound closure was severely debilitated by peptide treatment. Thus, the IC50 of the treatment peptide resulted in obstruction of cellular migration and proliferation (specifically seen in SKOV3, where KI67 expression decreased with treatment). However, the gene expression profile of SNU499 cells tested for EMT and autophagic genes did not change upon treatment, suggesting possible changes in the epigenetic or protein levels [25,26,27]. Contrary to that, SKOV3 treated with the peptide showed reduced expression of proliferation marker KI67 and adhesion marker B-catenin, while also exhibiting an increase in EMT marker Vimentin and autophagic markers ATG5 and ATG6, with no expression of ATG7. This may indicate that exposure of SKOV3 cells to the peptide reduced gene expression associated with proliferation, cell adhesion, and autophagy while also displaying increases in EMT-associated marker Vimentin [28]. Further study of EMT markers and protein expression levels is suggested for a more comprehensive understanding of the implications of the treatment on cancer cells.

To verify the possible cell death mechanism, we used Annexin V/PI staining. We observed that there was a clear bias towards early apoptosis for peptide treated SNU449 cells as opposed to the control. Apoptosis was characterized by morphological changes, nucleic acid fragmentation, and loss of membrane asymmetry [29]. Based on our Annexin V findings, apoptosis was shown to be the dominant resulting mode of cell death caused by the peptide treatment. In treated cells, 51% of cells were in early apoptosis in comparison to untreated control that showed only 6% early apoptotic death. In addition to this, 18% of the peptide-treated cells were in late-stage apoptotic/ necrotic. These results suggest that apoptosis is the most pronounced cellular death upon treatment [30].

The targeted cytotoxicity of the peptide and its ability to penetrate or disrupt the cellular membrane of human erythrocytes was further evaluated through hemolysis assay. This was determined by measuring the OD of the released hemoglobin upon treatment, reflecting cellular rupturing and/or penetration [31]. Absorbance values of IC50 (76.4 µM) and IC25 (53.87 µM) values, from peptide treated SNU449 (the highest concentration from any cancer cell line), was negligible, showing a similar effect to that of saline, the baseline negative control. Thus we inferred that the peptide did not penetrate normal red blood cells, during the incubation period. These results suggest the peptide’s cytotoxicity was higher on relatively anionic cancer cells while sparing normal neutral cells (erythrocytes). We thus would suggest intravenous induction as a mode of administration for potential in vivo examination. Additionally, the 22.1 min blood-borne predicted half-life of the peptide is an indicator of its ability to persist within the bloodstream, which is similar to the 20 - 30 min half-life of platinum-based drug cisplatin [32]. The promising anticancer properties of our peptide established on cell viability, proliferation, morphology, and migration and its minimal hemolytic activity, promote its novel therapeutic potential as an anticancer drug [33].

To further characterize our novel 37-mer peptide, we sought to establish its antimicrobial activity on both gram-negative and gram-positive bacterial strains. Induction of 118.7 µM and 161.6 µM of the peptide resulted in a comparable antibacterial effect on *S. aureus* and *E. coli* at the 24 h time point. Peptide treatment showed a significant reduction in colony formation for *E. coli* at 118.7 µM and 161.6 µM. Antimicrobial peptides are known to be part of the natural Host defense mechanism against pathogens (Host Defense Peptides) [34] in the innate immune system of all species [35]. Most of the studied AMPs possess a broad-spectrum effect on all microbes, including bacteria, viruses, and fungi [9]. It is noteworthy that *S. aureus* and *E. coli* are part of the human normal flora. *S. aureus* colonizes the skin and most of the mucosal membranes’ microbiota and E. coli colonizes the gastrointestinal tract [36, 37]. Consequently, the antimicrobial effect of the peptide suggests an effect of the peptide upon administration on the normal human microbiota. However, the IC50 of the peptide established on all tested cell lines is much lower than the tested peptide concentrations on both bacterial strains, suggesting a marginal effect of the peptide on the normal flora upon its administration as a therapeutic agent. The negligible hemolytic activity of the peptide on normal erythrocytes suggests that intravenous administration is preferable.

The antimicrobial and the anticancer properties of the novel peptide have been attempted to be established throughout this current work. This dual property has the potential to be utilized as a promising anticancer therapeutic agent, with rapid and targeted action, hindering cancerous cells from acquiring metastatic properties [9].

## Conclusion

Peptide drug approaches are a novel exploratory therapeutic for cancer research, with the potential to overcome the existing issues of present therapies. Our novel antimicrobial peptide has displayed the capacity to have a preliminary anticancer effect on hepatocellular carcinoma cell lines HepG2 and SNU449, and an ovarian cancer cell line SKOV3, while showing a moderate effect on immortalized mammalian fibroblasts. Morphologically degenerated cells, exposed to the peptide, were hindered during cell migration, and underwent early apoptotic cell death while showing limited cytotoxic effects on erythrocytes. The peptide’s electrostatic and structural properties significantly increased its anticancer profile. The dual properties of the peptide against cancer cells and bacterial strains supported its novelty. Thus, in furthering the validation of this work, we recommend the expansion to test our anticancer peptide on a cancer animal model or more physiologically relevant 3D tumor models.

## Supporting Information

**S1 Table. RT-PCR primer parameters: sequences, annealing temperature, cycle number, and amplicon size.**

**S1 Fig. Effect of 37-mer peptide treatment on HeLa cells over 24 h. (A)** Data showed negligible effect on cell viability with an increase in ACP concentration (“*” denotes P<0.01, “**” denotes P<0.001, n=3). **(B)** Treatment of HeLa cells with 121.5 µM. Untreated cells formed compact circular condensed attached cells. Peptide treatment caused cell morphology to become more rounded, sparse, and detach from the plate. Highlighted circle regions showed rounded dead cells releasing cellular vacuoles.

**S2 Table. Summary of IC50, hill coefficient, and R2 values obtained for peptide treatment of HeLa and MCF7 cell lines.**

**S2 Fig. Gene expression of SNU449 cells of certain Epithelial to Mesenchymal markers and Autophagy genes was not affected by peptide treatment.** (A) Gene expression level of *KI67, B-Catenin, Survivin, Bax, N-cadherin, E-cadherin, and Vimentin* from SNU449 24 h treated (IC50) and untreated cells (n=3). Gene expression data were generated and normalized against *GAPDH* as a control. Expression data did not statistically change between untreated and treated cells. (B) Autophagy gene expression of SNU449 cells from *ATG5, ATG6, and ATG7* was determined. Treated cells did not display a statically significant variation in expression in epithelial to mesenchymal transition genes and autophagy genes.

**S3 Fig. 37-mer peptide treatment affected gene expression levels of certain EMT and Autophagy markers of SKOV3 cells. (A)** SKOV3s were treated with peptide IC50 for 24 h and profiled for EMT and Autophagy markers. Gene expression data were generated and normalized against *GAPDH* as a control. *KI67, B-Catenin*, and *Vimentin* differential expression were significant in treated cells (*KI67* and *B-Catenin* decreased, while *Vimentin* increased) compared to control. **(B)** Expression of Autophagy genes showed a significant increase of *ATG5* and *ATG6* treated cells, compared to untreated SKOV3s, while exposure to the peptide inhibited expression of *ATG7* compared to untreated cells (“**” denotes P<0.001, “****” denotes P<0.00001, n=3).

